# 3-hydroxykynurenine is a ROS-inducing cytotoxic tryptophan metabolite that disrupts the TCA cycle

**DOI:** 10.1101/2023.07.10.548411

**Authors:** Jane L. Buchanan, Adam J. Rauckhorst, Eric B. Taylor

## Abstract

Tryptophan is an essential amino acid that is extensively characterized as a regulator of cellular function through its metabolism by indoleamine 2,3-deoxygenase (IDO) into the kynurenine pathway. However, despite decades of research on tryptophan metabolism, the metabolic regulatory roles of it and its metabolites are not well understood. To address this, we performed an activity metabolomics screen of tryptophan and most of its known metabolites in cell culture. We discovered that treatment of human colon cancer cells (HCT116) with 3-hydroxykynurenine (3-HK), a metabolite of kynurenine, potently disrupted TCA cycle function. Citrate and aconitate levels were increased, while isocitrate and all downstream TCA metabolites were decreased, suggesting decreased aconitase function. We hypothesized that 3HK or one of its metabolites increased reactive oxygen species (ROS) and inhibited aconitase activity. Accordingly, we observed almost complete depletion of reduced glutathione and a decrease in total glutathione levels. We observed a dose-dependent decrease in cell viability after 48 hours of 3HK treatment. These data suggest that raising the intracellular levels of 3HK could be sufficient to induce ROS-mediated apoptosis. We modulated the intracellular levels of 3HK by combined induction of IDO and knockdown of kynureninase (KYNU) in HCT116 cells. Cell viability decreased significantly after 48 hours of KYNU knockdown compared to controls, which was accompanied by increased ROS production and Annexin V staining revealing apoptosis. Finally, we identify xanthommatin production from 3-HK as a candidate radical-producing, cytotoxic mechanism. Our work indicates that KYNU may be a target for disrupting tryptophan metabolism. Interestingly, many cancers exhibit overexpression of IDO, providing a cancer-specific metabolic vulnerability that could be exploited by KYNU inhibition.

## Introduction

Tryptophan is an essential amino acid that is used for protein synthesis, neurotransmitter production, and de novo NAD+ synthesis via the kynurenine pathway. Almost 95% of available tryptophan is directed through the kynurenine pathway into metabolites that can exhibit immunosuppressive properties, neuroprotective properties, or neurotoxic effects (1–3). In cancer, kynurenine has a potent immunosuppressive effect on the tumor microenvironment. Tumor overexpression of indoleamine 2,3-deoxygenase (IDO), the rate-limiting enzyme of the kynurenine pathway, is associated with increased levels of kynurenine, which can bind to the aryl hydrocarbon receptor and decrease T cell effector function and proliferation (4). In neurodegenerative diseases, kynurenine pathway metabolites that are downstream of kynurenine have been shown to have either a neuroprotective effect (kynurenic acid) or neurotoxic effect (quinolinic acid). Quinolinic acid can stimulate *N*-methyl-D-aspartate (NMDA) receptors leading to excitatory neuron death (5), while kynurenic acid acts as an antagonist for ionotropic glutamate receptors, and thus protects against excitotoxicity (6). Other kynurenine pathway metabolites, such as 3-hydroxykynurenine (3HK) have also been implicated in neuronal toxicity. At low levels, 3HK functions as an antioxidant and can scavenge reactive oxygen species (ROS) such as peroxyl radicals (7, 8). At high levels or under oxidative conditions, 3HK can produce ROS at concentrations that induce apoptosis and neuronal death (9–11). Increased levels of 3HK in the brain have been associated with several neurodegenerative diseases, including Huntington’s Disease (HD) (12, 13). Interestingly, in a transgenic mouse model of HD, 3HK was the only kynurenine pathway metabolite to increase following an intrastriatal injection of kynurenine (14). Furthermore, brain tissue from these mice exhibited significantly lower activity of kynureninase, the enzyme responsible for metabolizing 3HK in mammals.

Overall, the tryptophan metabolite literature can be summarized by binning metabolites into two main disease contexts: immunosuppression that may contribute to tumor progression (kynurenine), and neurotrophic or neurotoxic (kynurenic acid, quinolinic acid, 3HK). However, it is unclear whether tryptophan metabolites that cause neuronal toxicity could also exert a cytotoxic effect on cancer cells. Given that 3HK is thought to induce neuronal apoptosis via ROS and not through excitotoxicity mechanisms, 3HK could potentially exert cytotoxic effects against cancer cells, especially in IDO-overexpressing tumors that exhibit high tryptophan consumption.

In this study, we performed an activity metabolomics screen in cancer cells with tryptophan metabolites and found that 3HK was the most potent disruptor of TCA cycle metabolism and glutathione levels. Induction of IDO expression, followed by kynureninase knockdown, led to significantly increased levels of ROS, apoptosis, and cell death compared to the non-targeting control. Thus, these results suggest that kynureninase inhibition may be an effective way to disrupt tryptophan metabolism and induce cell death in IDO-expressing cancer cells.

## Methods

### Custom Cell Culture Media (CM)

Ingredients were bought or made in bulk and frozen in aliquots at -20°C (**Supplemental Table 1**). One day prior to cell culture, reagents were thawed in a 37°C water bath and combined. pH was adjusted to 7.4 +/-0.05, and media was filtered using a 0.22 µm sterile bottle-top filter. Media was kept at 4°C until time of use. Due to its sensitivity to degradation, glutamine was not added to media until time of use.

### Crystal Violet Cell Cytotoxicity Assay

HCT15 cells were plated in 1:1 DMEM:CM on a clear 96-well plate. After 18 hours, cells were treated with 200 µl of CM (CTRL) or CM plus an amino acid (2 mM). Cells were then grown for 48 hours, followed by a live/dead assay using the Crystal Violet Cell Cytotoxicity Assay Kit (BioVision, #K329) according to the manufacturer’s directions. Briefly, cells were washed with 200 µl of 1x washing solution and incubated in 50 µl of crystal violet staining solution for 20 minutes. Cells were washed four times with 200 µl of 1x washing solution to remove dead cells. Cells were then incubated in 100 µl of solubilization solution for 20 minutes, and absorbance was measured at 595 nm.

### Cell-TiterGlo Luminescent Cell Viability Assay

HT29, 293T, U2OS, and 3T3-L1 cells were plated on an opaque 96-well plate in DMEM. After 18 hours, cells were treated with 100 µl of DMEM (vehicle) or DMEM plus 2 mM tryptophan for 24 hours (Figure 1B). For 3HK dose treatment cells were plated in 1:1 DMEM:CM, the following day culture media was changed to 100 µl of CM (vehicle) or CM plus 3HK in a 2-fold dilution series for 48 hours (Figure 3F). For IFN-γ + KNYU-knockdown experiments, 24 hours after DsiRNA transfection (detailed below), cells were treated with 50 ng/µl IFN-γ in 1:1 DMEM:CM. The following day media was changed to CM (Figure 4). At the appropriate time point, 100 µl of CellTiter-Glo reagent (Promega #G7570) was added to each well. The plate was gently mixed for 2 minutes to induce lysis, incubated at room-temperature for 10 minutes, and then luminescence was measured.

**Figure 1.**
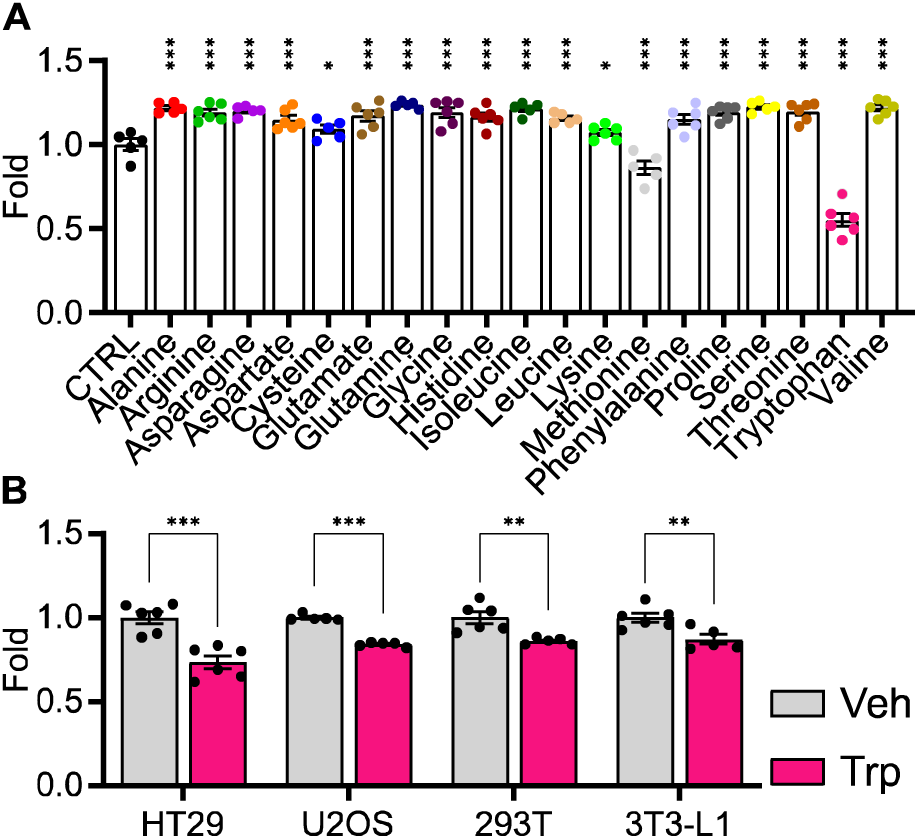
A) Bar graph showing the effect of individual amino acids (2 mM) on HCT15 cell proliferation as measured by crystal violet assay after 48 hours of treatment (n = 5-6, statistical analysis by one-way ANOVA with Holm-Sidak post-hock comparison compared to control (CTRL)). B) Bar graph showing the effect of tryptophan (2 mM) on cell viability in four different cell lines (HT29, U2OS, 293T, 3T3-L1) as measured by CellTiter-Glo assay after 24 hours of treatment (n = 5-6, statistical analysis by Student’s t-test compared to Vehicle (Veh)). Data shown as mean ± SEM. * p < 0.05, ** p < 0.01, ***p < 0.001.

**Figure 2.**
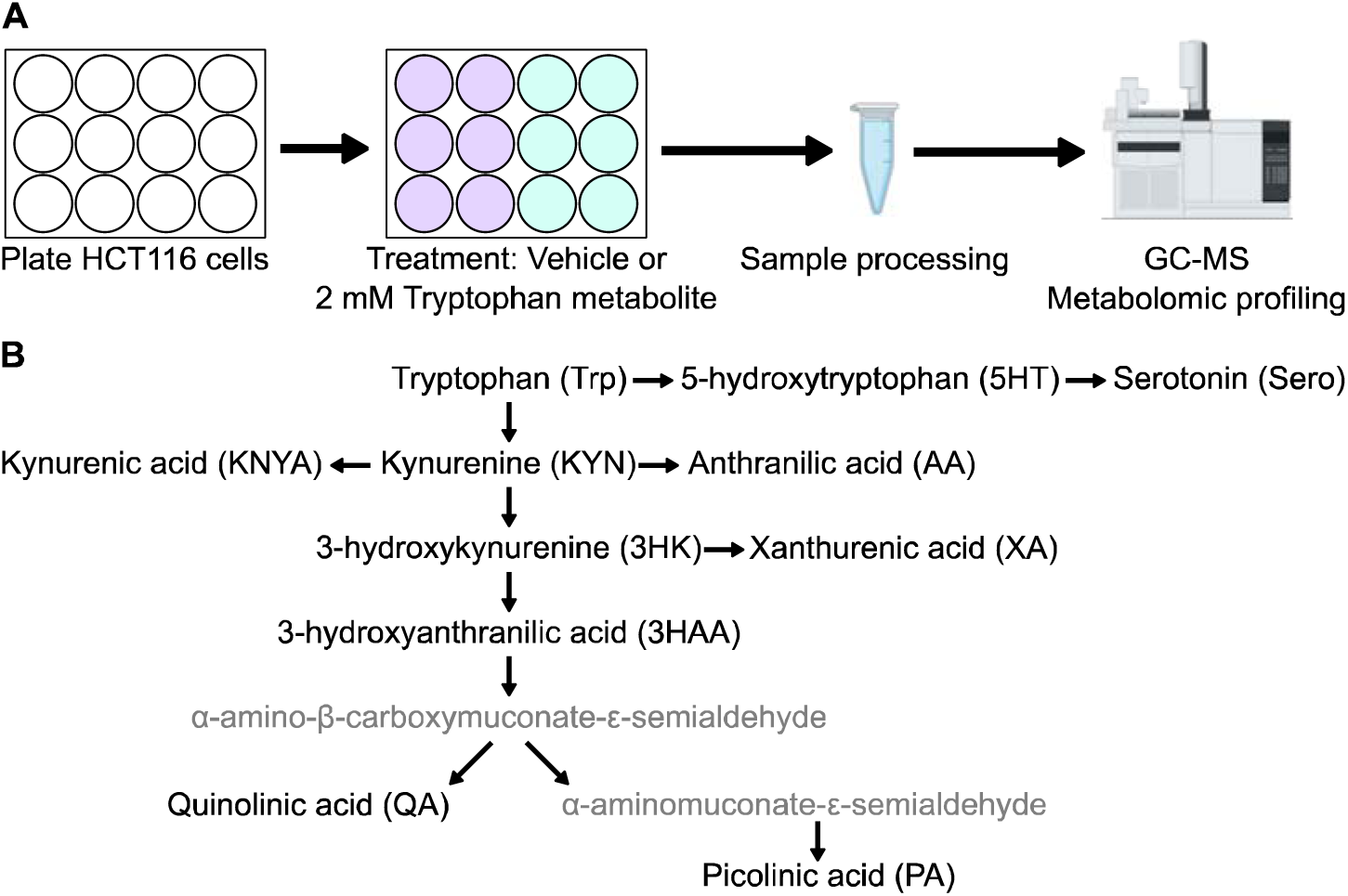
A) Experimental design for metabolic profiling. HCT116 cells were treated with tryptophan metabolites and analyzed via gas chromatography mass spectrometry (GC-MS). B) Diagram of tryptophan metabolism and metabolites tested in the experiment (black font, abbreviations shown in parentheses).

**Figure 3.**
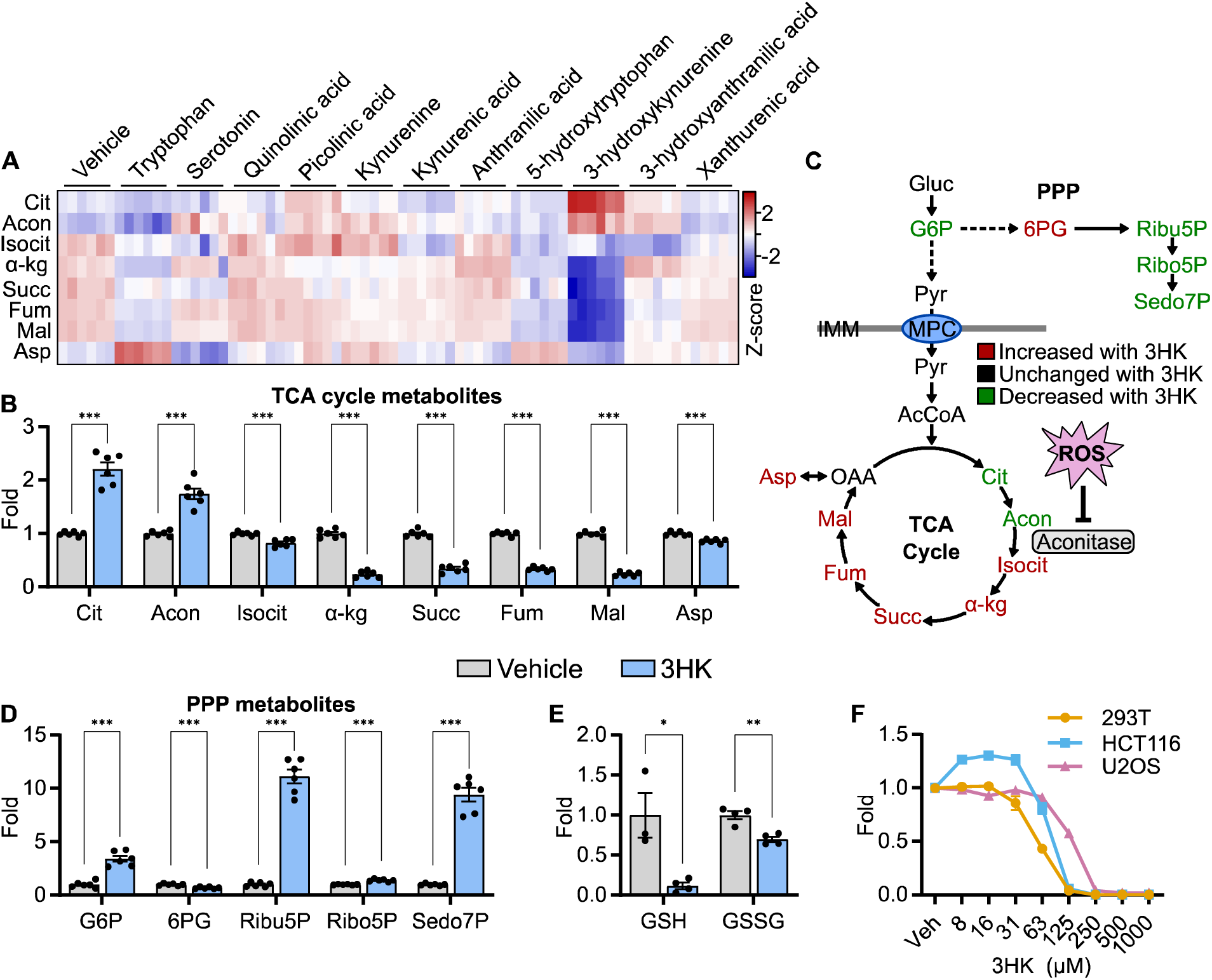
A) Heat map showing the Log2 fold change of TCA cycle intermediates in HCT116 cells following tryptophan metabolite treatment for 6 hours (n = 6). B) Bar graphs showing relative fold changes of TCA cycle intermediates in HCT116 cells treated with 3HK (2 mM) for 6 hours (n = 6, statistical analysis by Student’s t-test compared to Vehicle). C) Schematic showing TCA cycle and pentose phosphate pathway (PPP) metabolites color-coded based on their direction of change following treatment with 3HK (2 mM) for 6 hours. D) Relative fold changes of PPP intermediates in HCT116 cells treated with 3HK (2 mM) for 6 hours (n = 6, statistical analysis by Student’s t-test compared to Vehicle). E) Bar graphs showing reduced (GSH) and oxidized glutathione (GSSG) in HCT116 cells treated with 3HK (2 mM) for 6 hours. (n = 3-4, statistical analysis by Student’s t-test compared to Vehicle). F) Line graph showing cell viability in three cell lines (293T, HCT116, U2OS) treated with 3HK at the concentrations indicated after 48 hours of treatment (n = 3-4). Data shown as mean ± SEM. * p < 0.05, ** p < 0.01, ***p < 0.001. Abbreviations: Gluc, glucose; G6P, glucose 6-phosphate; Ribu5P, ribulose 5-phosphate; Ribo5P, ribose 5-phosphate; Sedo7P, sedoheptulose 7-phosphate; Pyr, pyruvate; AcCoA, acetyl-CoA; Cit, citrate; Acon, aconitate; Isocit, isocitrate; α-kg, α-ketoglutarate; Succ, succinate; Fum, fumarate; Mal, malate; OAA, oxaloacetate; Asp, aspartate; ROS, reactive oxygen species; IMM, inner mitochondrial membrane.

**Figure 4.**
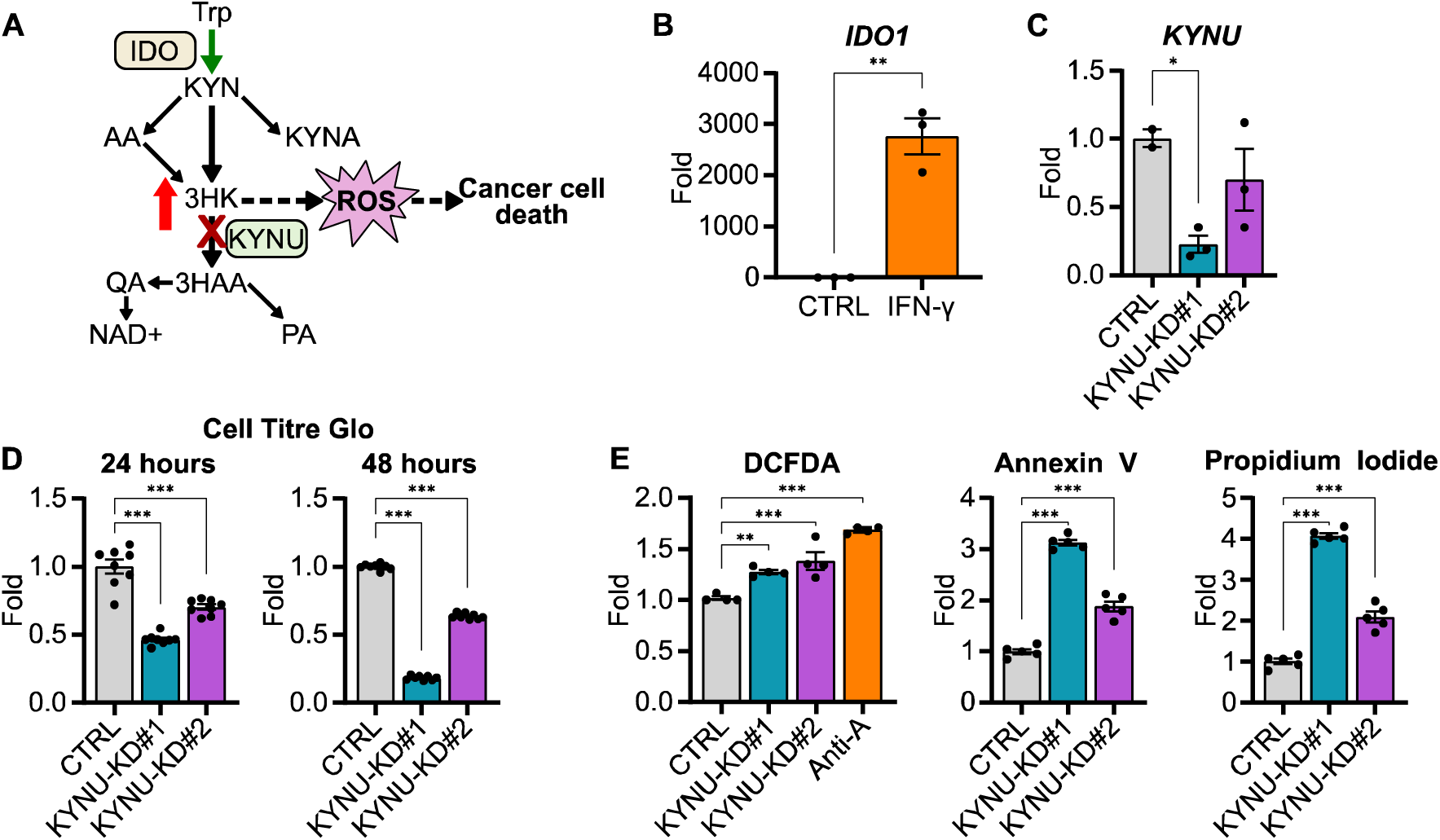
A) Schematic showing the proposed mechanism for increasing 3HK in cancer cells to mediate their death. B) Bar graph showing *IDO1* mRNA levels in HCT116 cells following treatment of IFN-γ (50 ng/µL) for 24 hours. (n = 3, statistical analysis by Students t-test compared to control (CTRL)). C) Bar graph showing *KYNU* mRNA levels in HCT116 cells following treatment of following DsiRNA mediated KYNU knockdown (KYNU-KD). (n = 2-3, statistical analysis by one-way ANOVA with Holm-Sidak post-hock comparison compared to non-targeting control (CTRL)). D) Bar graphs showing viability of HCT116 cells after 24 (left) and 48 (right) hours of KYNU knockdown compared to non-targeting control (n = 8, statistical analysis by one-way ANOVA with Holm-Sidak post-hock comparison compared to non-targeting control (CTRL)). E) Bar graphs showing DCFDA (left), Annexin V (center), and propidium iodide (right) staining in HCT116 cells after 48 hours of KYNU knockdown (n = 3-5, statistical analysis by one-way ANOVA with Holm-Sidak post-hock comparison compared to non-targeting control (CTRL)). Data shown as mean ± SEM. * p < 0.05, ** p < 0.01, ***p < 0.001.

### siRNA Transfection

Predesigned DsiRNAs targeting KYNU (Design ID: hs.Ri.KYNU.13.2, hs.Ri.KYNU.13.3), a HPRT-S1 positive control DsiRNA, and a negative control DsiRNA were obtained from Integrated DNA Technologies. DsiRNAs were resuspended in nuclease-free duplex buffer. For each well of a 96-well plate, 6 pmol of DsiRNA were diluted in 100 µL Opti-MEM media (Gibco #31985062) and pipette-mixed. 0.3 µL/well of Lipofectamine RNAiMAX (Invitrogen #13778075) was added, pipette-mixed, and incubated at room-temperature for 20 minutes. 20 µL of diluted HCT116 cells were added to each well to achieve 30-50% confluence after 24 hours. Final concentration of RNA was 10 nM. Cells were grown at 37°C in a CO_2_ incubator until time of cell viability assay, knockdown assay, or flow cytometry.

### Quantitative PCR (qPCR)

Total RNA from HCT116 cells was extracted using the TRIzol (Invitrogen #15596026) according to the manufacturer’s directions. cDNA synthesis of equal amounts of RNA from each sample was achieved using the Applied Biosystems High-Capacity cDNA Reverse Transcription Kit (Thermo Fisher #4368814). SYBR Green ER SuperMix (Thermo Fisher #11762500) was used during qPCR. Relative abundance of mRNA was normalized to the abundance of cyclophilin mRNA. Primer sequences (5’-3’): KYNU-1F: 5;GGGGGTGCCAGCTAACAATA, KYNU-1R: TCCGCTTGTCACAAACCACT, KYNU-2F: GATCCTGTTCAGTGGGGTGC, KYNU-2R: GCCAACATAACAACCCTTCGC; Cyclophilin-F: GGAGATGGCACAGGAGGAAA, Cyclophilin-R: GCCCGTAGTGCTTCAGTTT.

### Annexin V and Propidium Iodide for Apoptosis and Live/Dead Staining

Cells were dissociated using Accutase (Invitrogen #00-4555-56) (Trypsin) for 20 minutes at 37°C. The single cell suspension was then washed twice in PBS -/-. 5 µL of Annexin V-Alexa488 labeling buffer was added to 100 µL of cells for 15 minutes. 5 µL of propidium iodide (PI) buffer was then added for 15 minutes. Cells were immediately analyzed using a 4-laser acoustic-focusing Attune NxT Flow Cytometer (ThermoFisher). Signal was read at an Ex/Em of 490/525 nm for Annexin V and 535/617 nm for PI. Data were analyzed using FlowjoX.

### DCFDA Cellular ROS Detection Assay

Cells were dissociated using Accutase (Trypsin) for 20 minutes at 37°C. The single cell suspension was then washed twice in PBS -/-. Cells were stained and resuspended with 2′,7′-dichlorofluorescin diacetate (DCFDA) (20 µM) for 30 minutes. Cells were immediately analyzed using a 4-laser acoustic-focusing Attune NxT Flow Cytometer (ThermoFisher). Signal was read at an Ex/Em of 485/535 nm. Data were analyzed using FlowjoX.

### Xanthommatin synthesis

Xanthommatin was prepared as previously described (15). Briefly, 2 mM 3HK was mixed with 2 mM H_2_O_2_, and 10 pg/ml HRP in sodium phosphate buffer (50 mM, pH 7.4). The reaction was carried out at 20°C with exposure to room air.

### Metabolomics Sample Preparation

6-well plates were treated with 2 mM of the indicated tryptophan metabolite in CM. After 6 hours of treatment, media was quickly removed, and plates were washed 2x with ice-cold PBS and 2x with ice-cold water before being snap frozen with liquid nitrogen. Cell plates were lyophilized overnight and then metabolites were extracted. To extract metabolites, 1 ml of an extraction buffer composed of 2:2:1 methanol/acetonitrile/water containing internal standards at 1 μg/ml each (D4-Citric Acid, 13C5-Glutamine, 13C5-Glutamic Acid, 13C6-Lysine, 13C5-Methionine, 13C3-Serine, D4-Succinic Acid, 13C11-Tryptophan, and D8-Valine; Cambridge Isotope Laboratories) was added to the lyophilized cell samples. Cells were scraped for 20 seconds and collected into microcentrifuge tubes, flash-frozen in liquid nitrogen, and thawed in a room temperature water bath sonicator for 10 minutes. Next, samples were rotated at –20°C for 1 hour and then centrifuged at 4°C for 10 minutes at 21,000 × g. 150 µL of the cleared metabolite extracts were transferred to glass autosampler vials for derivatization and GC-MS analysis and 300 µL of the cleared metabolite extracts were transferred to fresh microcentrifuge tubes for LC-MS analysis. An equal volume of each extract was pooled to serve as a quality control (QC) sample, which was analyzed at the beginning, end, and at a regular interval throughout the analytical run. Extraction buffer alone was analyzed as a processing blank sample. Metabolite extracts, the quality control sample, and the processing blank were evaporated to dryness using a speed-vacuum.

### GC-MS Analysis

The dried samples, QC sample, and processing blank sample, were derivatized using methoxyamine hydrochloride (MOX) and N,O-Bis(trimethylsilyl)trifluoroacetamide (TMS) (Sigma). Briefly, dried extracts were reconstituted in 30 μL of 11.4 mg/ml MOX in anhydrous pyridine (VWR), vortexed for 10 minutes, and heated at 60°C for 1 hour. Next, 20 μL TMS was added to each reconstituted extract, vortexed for 1 minute, and heated at 60°C for 30 min. The derivatized samples, QC samples and processing blank samples were immediately analyzed using GC/MS.

GC chromatographic separation was conducted on a Thermo Trace 1300 GC with a TraceGold TG-5SilMS column (0.25 µM film thickness; 0.25mm ID; 30 m length). 1 μL of derivatized sample, QC, or blank was injected. The GC was operated in split mode with the following settings: 20:1 split ratio; split flow: 24 μL/min, purge flow: 5 mL/min, Carrier mode: Constant Flow, Carrier flow rate: 1.2 mL/min). The GC inlet temperature was 250°C. The GC oven temperature gradient was as follows: 80°C for 3 min, ramped at 20°C/min to a maximum temperature of 280°C, which was held for 8 min. The injection syringe was washed 3 times with pyridine between each sample. Metabolites were detected using a Thermo ISQ single quadrupole mass spectrometer. The data was acquired from 3.90 to 21.00 min in EI mode (70eV) by single ion monitoring (SIM).

### LC-MS Analysis

Dried extracts were reconstituted in 30 μL of acetonitrile/water (1:1, V/V), vortexed for 10 minutes, and incubated at –20°C for 18 hours. Samples were then centrifuged at 4°C for 2 minutes at 21,000 × g and the supernatant was transferred to autosampler vials for analysis. For LC chromatographic separation, 2 µL of reconstituted samples, QC sample, and processing blank sample were separated on a Millipore SeQuant ZIC-pHILIC (2.1 × 150 mm, 5 µm particle size, Millipore Sigma #150460) column with a ZIC-pHILIC guard column (20 × 2.1 mm, Millipore Sigma #150437) attached to a Thermo Vanquish Flex UHPLC. The mobile phase comprised Buffer A [20 mM (NH_4_)_2_CO_3_, 0.1% NH_4_OH (v/v)] and Buffer B [acetonitrile]. The chromatographic gradient was run at a flow rate of 0.150 mL/min as follows: 0–20 min-linear gradient from 80 to 20% Buffer B; 20-20.5 min-linear gradient from 20 to 80% Buffer B; and 20.5–28 min-hold at 80% Buffer B. Data were acquired using a Thermo Q Exactive MS operated in full-scan, polarity-switching mode with a spray voltage set to 3.0 kV, the heated capillary held at 275°C, and the HESI probe held at 350°C. The sheath gas flow was set to 40 units, the auxiliary gas flow was set to 15 units, and the sweep gas flow was set to 1 unit. MS data acquisition was performed with polarity switching in a range of m/z 70–1,000, with the resolution set at 70,000, the AGC target at 10e6, and the maximum injection time at 200 ms.

### Mass Spectrometry and Data Analysis

Acquired LC-MS and GC-MS data were processed using the Thermo Scientific TraceFinder 5.0 software. Targeted metabolites were identified by matching one target and at least one confirming ion and retention time (GC-MS) or accurate mass and retention times (LC-MS) to the University of Iowa Metabolomics Core Facility’s in-house library of confirmed standards. After peak area integration by TraceFinder, NOREVA software was applied for signal drift correction on a metabolite-to-metabolite basis using the pooled QC sample that had been analyzed throughout the instrument run (16). Within individual samples, Post-NOREVA metabolite levels were divided by the total metabolite load (sum of all metabolite values) to equally weight individual metabolites for comparison across samples.

## Results

### Tryptophan decreases cell proliferation in colon cancer cells

An amino acid screen of all 20 essential and non-essential amino acids was performed to assess the differential effects of amino acids on cancer cell proliferation. Because commercially available cell culture media such as DMEM or RPMI contain variable concentrations of amino acids, we designed a chemically defined, custom minimal media containing all amino acids present at 0.3 mM, referred to as CM. HCT15 cells, a colon cancer cell line, were plated in 50:50 DMEM:CM and left to attach overnight. Media was then changed to 100% CM, and cells were treated with 2 mM of individual amino acids. After 48 hours, cell proliferation was measured by crystal violet dye, which binds to ribose-like molecules such as DNA and assumes dead cells become unattached and are washed away. Most amino acids increased cell proliferation compared to CM only, however, cells treated with tryptophan exhibited nearly a 50% decrease in cell viability (**Fig. 1A**). This suggests that, compared to other amino acids, tryptophan is uniquely cytotoxic to HCT15 cells at high concentrations.

To test whether tryptophan inhibited the growth of other cell lines and persisted in conventional cell culture media, two human cancer cell lines (HT29, U2OS), a transformed human embryonic kidney cell line (293T), and a mouse, fibroblast-like cell line (3T3-L1) were grown in DMEM for 24 hours and then treated with 2 mM tryptophan or vehicle. After 24 hours, cells were assayed by CellTiter-Glo, which determines cell viability by measuring ATP levels. All cell lines showed significant decreases in cell viability compared to vehicle (**Fig. 1B**). These data demonstrate that the cytotoxic effects of high concentration tryptophan are not exclusive to HCT15 cells or an artifact of the CM.

### 3HK potently disrupts TCA cycle metabolism, glutathione levels, and cell viability

To better understand the metabolic effects that could be contributing to decreased cell viability, an activity metabolomics screen was performed in HCT116 cells with tryptophan and its metabolites (**Fig. 2A**). Cells were treated with 2 mM tryptophan metabolites (**Fig. 2B**) or vehicle for 6 hours and metabolic profiling was performed by GC-MS. Because prior pilot studies showed evidence of tryptophan affecting TCA cycle metabolites, we examined the effects of tryptophan metabolites on the TCA cycle first and found that 3HK exhibited more potent disruption than the other tryptophan metabolites (**Fig. 3A**). Interestingly, proximal TCA cycle metabolites such as citrate and aconitate were significantly increased, while distal TCA cycle metabolites were drastically and significantly decreased, demonstrating a clear flux disruption between aconitate and isocitrate (**Fig. 3B**). Because aconitase, the enzyme that converts aconitate to isocitrate, contains an iron-sulfur cluster that is sensitive to oxidation, we hypothesized that ROS from 3HK or one its products could be inhibiting aconitase activity and preventing metabolism of citrate and aconitate (**Fig. 3C**). Significantly increased levels of glucose 6-phosphate, ribose 5-phosphate, and ribulose 5-phosphate indicated increased flux of carbon through the pentose phosphate shunt, which is used to produce NADPH for the reduction of the antioxidant glutathione (**Fig. 3D**). Accordingly, metabolic profiling of the same cell samples on LC-MS revealed significantly decreased levels of reduced and oxidized glutathione (**Fig. 3E**), which is a metabolic signature of ROS. Next, we then assessed the dose-dependent effects of 3HK on cell viability (293T, U2OS, HCT116). All cell lines tested exhibited a dose-dependent decrease in cell viability as measured by CellTiter-Glo (**Fig. 3F**). Together, these data demonstrate that 3HK potently affects normal cellular metabolism and is likely responsible for the cytotoxic effects of high tryptophan concentrations.

### Induction of IDO followed by kynureninase knockdown increases ROS, apoptosis, and cell death in colon cancer cells

Given the detrimental effects of 3HK on cell metabolism and viability, we hypothesized that 3HK accumulation resulting from kynureninase inhibition in IDO overexpressing cancer could result in cell toxicity and death (**Fig. 4A**). To test this hypothesis, we induced IDO expression in HCT116 cells using the known IDO potentiator interferon-γ (IFN-γ) (**Fig. 4B**). Then, we utilized DsiRNAs to knocked down KYNU, the gene that encodes kynureninase, in IDO-induced cells and assessed cell viability at 24 and 48 hours (**Fig. 4C**). KYNU-knockdown decreased cell viability at both time points compared to the non-targeting control (**Fig. 4D**). Next, we fluorescently probed ROS, apoptosis, and cell death using DCFDA, Annexin V, and propidium iodide, respectively. Indeed, KYNU-knockdown increased ROS, apoptosis, and cell death in a dose dependent manner with the potency of KYNU-DsiRNA (**Fig. 4E**). This suggests that increasing 3HK levels in IDO overexpressing cancers may be sufficient to induce cell death.

### The ROS-generating 3HK metabolite xanthommatin increases with tryptophan administration

Though there is a clear association between 3HK and elevated cellular ROS, we aimed to better understand whether 3HK or one of its metabolites was producing the radical species. 3HK can spontaneously oxidize (or autoxidize) to form the radical-producing compound xanthommatin, a dimer of 3HK (15, 17). Interestingly, xanthommatin has been well-characterized in insect eye pigmentation, however, only one study mentions xanthommatin being produced by mammalian biological material (18). This studied showed that cytochrome c and cytochrome oxidase were able to convert 3HK to xanthommatin in rat liver mitochondria (18). Using a previously described xanthommatin synthesis method (15), we generated xanthommatin from 3HK (**Fig. 5A**). Xanthommatin analyzed by LC-MS/MS matched publicly available and published (19) xanthommatin fragmentation data, confirming the compound’s identity (**Fig. 5B**). We treated HCT116 cells with 0.2 or 2 mM 3HK for 6 hours and observed a ∼70-fold increase of xanthommatin at the 2 mM dose (**Fig. 5C**). Finally, we treated HCT116 cells with IFN-γ for 24 hours to induce IDO expression and then treated with tryptophan (1 mM) for 6 hours. Metabolic profiling revealed that xanthommatin was the only tryptophan metabolite to significantly increase (**Fig. 5D**). These data demonstrate that xathommatin is formed in HCT116 cells treated with tryptophan.

**Figure 5.**
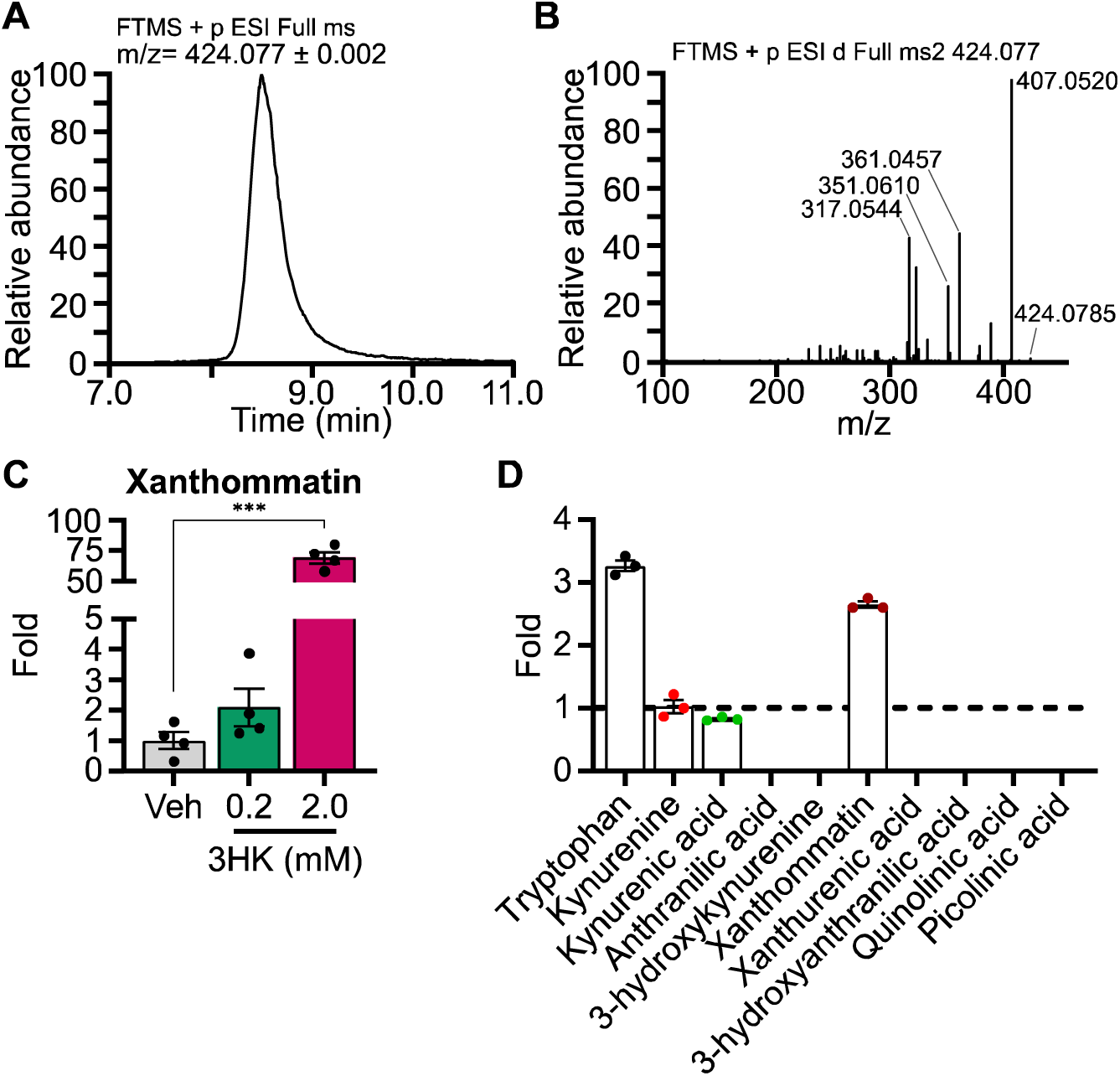
A) Line graph showing the extracted ion chromatogram of xanthommatin [M+H] molecular ion (424.077 m/z) viewed with a 2 milli mass unit tolerance. B) Line graph showing the xanthommatin molecular ion MS/MS mass spectra collected with a higher-energy collision dissociation (HCD) energy of 40%. C) Bar graph showing the relative fold xanthommatin levels in HCT116 cells treated with 3HK at the specified dosages for 6 hours (n = 4, statistical analysis by one-way ANOVA with Holm-Sidak post-hock comparison compared to vehicle treatment (Veh)). D) Bar graph showing relative fold xanthommatin levels in HCT116 treated with IFN-γ (50 ng/µL) for 24 hours and subsequent 1 mM tryptophan treatment for 6 hours compared to vehicle (dotted line). (n = 3, nd = not detected). Data shown as mean ± SEM. ***p < 0.001.

## Discussion

Tryptophan metabolites have been extensively studied, particularly in the context of tumor-mediated immunosuppression and neuronal protection or cytotoxicity (1–3). However, the metabolic effects of tryptophan metabolites have not been systematically assessed in a side-by-side comparison in cells before. By adding the same concentration of each tryptophan metabolite to cancer cells in a chemically defined, custom media and performing an activity metabolomics assay, we were able to show that 3HK exerts the greatest metabolic disruption on TCA cycle and pentose phosphate pathway metabolites. Increased levels of proximal TCA cycle metabolites and decreased distal TCA cycle metabolites after 3HK treatment suggest a ROS-mediated inhibition of aconitase activity. Furthermore, increased levels of pentose phosphate pathway metabolites and decreased glutathione levels are a metabolic signature of excess ROS. 3HK also decreased cell viability in multiple cell lines in a dose-dependent manner.

Interestingly, 3HK has been the focus of relatively few tryptophan metabolite studies, with some reviews failing to mention it all. Studies that have investigated it were predominantly limited to a neuronal context. 3HK’s ROS-mediated toxicity in cancer cells in this study indicate a metabolic vulnerability that could potentially be exploited in IDO-positive tumors. Inducing IDO expression to increase tryptophan catabolism to 3HK and knocking down KYNU to decrease 3HK degradation were sufficient to induce significant apoptosis and cell death in culture. These results raise the possibility that inhibition of kynureninase in IDO-positive tumors could lead to sufficient intracellular levels of 3-hydroxykynurenine/xanthommatin to mediate cytotoxic effects. Notably, though small molecule inhibitors have been developed to target IDO and other kynurenine pathway enzymes, no inhibitors have been developed to target kynureninase.

The detection of xanthommatin in HCT116 cells treated with tryptophan is a novel finding that warrants future investigation. In vitro studies showed that autoxidation of 3HK or incubating 3HK with horseradish peroxidase and hydrogen peroxide, produced a radical that is detected when a xanthommatin peak appears, suggesting that xanthommatin is the source of the radical (15, 17). Xanthommatin has been well-studied in the context of insect eye pigmentation, where 3HK is enzymatically metabolized by cardinal, an insect-specific heme peroxidase (20). Interestingly, xanthommatin has never been reported in humans, despite the fact the top cardinal orthologs in humans are also heme peroxidases (https://flybase.org/reports/FBgn0263986.html). Given the number of peroxidases that are present in human cells and the fact that 3HK can also autoxidize, it is plausible that xanthommatin is not only produced in mammalian cells, as our data show, but that it could also be the source of ROS that are responsible for 3HK’s cytotoxicity. Future experiments could measure ROS production and cell viability when cells are treated with xanthommatin and assess whether xanthommatin is increased with KYNU inhibition. The discovery of xanthommatin in human cells prompts additional questions such as where is xanthommatin generated in mammalian cells? What enzymes in mammalian cells produce xanthommatin? What kind of physiological or disease conditions elicit xanthommatin production? What is xanthommatin’s role in mammalian cellular function?

Overall, our investigation of the effects of tryptophan metabolites on cellular metabolism revealed a novel metabolic vulnerability to high concentrations of 3HK, which could be exploited in cancer cells by inhibiting kynureninase. We also discovered that HCT116 cells produce xanthommatin, a 3HK metabolite that has not been described in human cells but has been shown to generate radical species in vitro. Furthermore, although 3HK was not detectable with tryptophan supplementation, xanthommatin levels were significantly increased, suggesting that xanthommatin from 3HK may catalyze ROS formation and mediate the cytotoxicity observed in our studies.

## Acknowledgements

This work was supported by grants NIH F30 DK127845 (JLB), ADA 1-18-PDF-060 and AHA CDA851976 (AJR), NIH R01 DK104998 (EBT) and the University of Iowa Healthcare Distinguished Scholars Award (EBT). We thank Kelly Falls-Hubert and Guillermo Romano Ibarra for technical assistance with fluorescent staining assays and flow cytometry experiments.

## Author Contributions

EBT and JLB conceived the study. EBT, JLB, and AJR designed the study. JLB performed experiments and collected data. JLB and EBT analyzed data. JLB, AJR, and EBT wrote, revised, and approved the manuscript. EBT supervised the study and AJR provided technical supervision.

---

**Supplemental Table 1:** Custom Cell Culture Media (CM)

| **Ingredients** | **Final Concentration** |
| --- | --- |
| **Amino Acids** | **mM** |
| L-glycine | 0.3 |
| L-alanine | 0.3 |
| L-arginine | 0.3 |
| L-asparagine | 0.3 |
| L-aspartic acid | 0.3 |
| L-cysteine | 0.3 |
| L-glutamic acid | 0.3 |
| L-glutamine | 0.3 |
| L-histidine | 0.3 |
| L-isoleucine | 0.3 |
| L-leucine | 0.3 |
| L-lysine | 0.3 |
| L-methionine | 0.3 |
| L-phenylalanine | 0.3 |
| L-proline | 0.3 |
| L-serine | 0.3 |
| L-threonine | 0.3 |
| L-tryptophan | 0.3 |
| L-tyrosine | 0.3 |
| L-valine | 0.3 |
| **Inorganic Salts** | **mM** |
| Calcium Chloride (CaCl<sub>2</sub> anhyd.) | 1.8018 |
| Magnesium Sulfate (MgSO<sub>4</sub> anhyd.) | 0.8139 |
| Potassium Chloride (KCl) | 5.3333 |
| Sodium Bicarbonate (NaHCO<sub>3</sub>) | 24 |
| Sodium Chloride (NaCl) | 110.3448 |
| Sodium Phosphate monobasic (NaH<sub>2</sub>PO<sub>4</sub>-H<sub>2</sub>O) | 0.9058 |
| **RPMI 1640 Vitamin Solution 100x** | **mM** |
| Biotin | 0.0008 |
| Ammonium chloride | 0.0400 |
| Choline chloride | 0.0214 |
| D-Calcium pantothenate | 0.0005 |
| Folic Acid | 0.0023 |
| Niacinamide | 0.0082 |
| Para-aminobenzoic acid | 0.0073 |
| Pyridoxine hydrochloride | 0.0049 |
| Riboflavin | 0.0005 |
| Thiamine hydrochloride | 0.0030 |
| Vitamin B12 | 0.000004 |
| i-Inositol | 0.1944 |
| **Other** | **mM** |
| D-Glucose | 5 |
| Sodium pyruvate | 0.1 |
| 3-Hydroxybutyric acid sodium salt | 0.05 |
| L-Carnitine hydrochloride | 0.04 |
| Sodium L-lactate | 0.9 |
| Gibco penicillin-streptomycin (10,000 U/mL) | 1x |
| Gibco dialyzed fetal bovine serum | 10x |

